# A single episode of sexual reproduction can prevent population extinction under multiple stressors

**DOI:** 10.1101/2024.04.26.589870

**Authors:** Yawako W. Kawaguchi, Masato Yamamichi

**Affiliations:** Center for Frontier Research, National Institute of Genetics, 1111 Yata, Mishima, Shizuoka 411-8540, Japan; Genetics Program, Graduate Institute for Advanced Studies, SOKENDAI, 1111 Yata, Mishima, Shizuoka 411-8540, Japan; School of the Environment, The University of Queensland, St. Lucia, Brisbane, Queensland 4072, Australia; Department of International Health and Medical Anthropology, Institute of Tropical Medicine, Nagasaki University, 1-12-4 Sakamoto, Nagasaki 852-8523, Japan; Institute for Multidisciplinary Sciences, Yokohama National University, 79-5 Tokiwadai, Hodogaya, Yokohama, Kanagawa 240-8501, Japan

**Keywords:** Adaptation, extinction, population biology, sex

## Abstract

Recent studies have shown that organisms can adapt to changing environments through rapid contemporary evolution. Although intraspecific genetic variation is necessary for rapid evolution to occur, little is known how genetic variation has been produced and maintained before rapid evolution. Here we show that a single episode of sexual reproduction can produce a large amount of intraspecific trait variation that allows population growth in degraded environments by laboratory experiments of a green alga, *Closterium peracerosum–strigosum–littorale* complex. We observed population dynamics of the alga under multiple stressors and confirmed that high salinity and low pH decreased population growth rates. By comparing parental and their hybrid F_1_ populations, we observed larger variation in population growth rates of F_1_ populations (i.e., transgressive segregation) when pH was low. Interestingly, even when parental populations had negative growth rates, some F_1_ populations showed positive growth rates in severe environmental conditions due to the large variation in population growth. By utilizing the recently obtained genomic information of the alga, we found greater enrichment of genes with copy number variations in terms related to pH stress than those to salt stress in a gene ontology (GO) enrichment analysis. Our results suggest that recombination and variation in the number of gene copies might produce large genetic variation in the F_1_ generation. This will be an important step toward a better understanding of evolutionary rescue, where rapid evolution prevents population extinction in changing environments.

## Introduction

Traditionally, evolutionary biologists tended to think that evolution is a slow process (Darwin 1859), but studies in the last quarter century have shown that organisms can flexibly adapt to changing environments through rapid contemporary evolution (Thompson 1998, Hairston et al. 2005, Schoener 2011, Hendry 2017, Sanderson et al. 2022). One of the most important implications of this phenomenon is that rapid evolution may prevent population extinction due to environmental changes, a process called “evolutionary rescue” (reviewed in Kinnison and Hairston 2007, Alexander et al. 2014, Carlson et al. 2014, Bell 2017). Previous studies have demonstrated that rapid evolution can prevent population extinction in severe environments by using mathematical modeling (e.g., Gomulkiewicz and Holt 1995, Orr and Unckless 2014, Uecker and Hermisson 2016, Orive et al. 2017), laboratory experiments (e.g., Bell and Gonzalez 2009, Lachapelle and Bell 2012, Lindsey et al. 2013), and field observations (e.g., Vander Wal et al. 2013, Isanta-Navarro et al. 2021).

It is well known, however, that evolution has no foresight and organisms cannot “prepare” for novel environmental changes in the future by maintaining genetic variation. How is genetic variation for rapid evolution produced and maintained in populations before the environmental changes? In addition to *de novo* mutations and immigration, classical studies in evolutionary biology have revealed that sexual reproduction is an important source for genetic variation (Williams 1975, Maynard Smith 1978, Bell 1982, Hartfield and Keightley 2012), and several experimental studies have indeed demonstrated that sexual reproduction can promote evolutionary rescue (Lachapelle and Bell 2012, Petkovic and Colegrave 2019, 2023). However, previous studies tended to compare experimental population dynamics of lineages with and without sexual reproduction and showed that sexual reproduction can increase the speed of adaptive evolution (Kaltz and Bell 2002, Goddard et al. 2005, Morran et al. 2009, Gray and Goddard 2012, Lachapelle and Bell 2012, Petkovic and Colegrave 2019, 2023). Thus, it was not clear how a single event of sexual reproduction can affect the amount of variation and the resultant rapid evolution. Colegrave et al. (2002) examined the effects of a single sexual episode on growth in various carbon sources, but they did not consider evolutionary rescue.

Here we show how a single episode of sexual reproduction can produce a large amount of trait variation and it makes population growth possible in degraded environments using laboratory experiments of a unicellular green alga, *Closterium peracerosum–strigosum–littorale* complex (Fig. 1A). The alga is a freshwater species belonging to the Zygnematophyceae, a group most closely related to land plants (Wickett et al. 2014). It usually grows vegetatively (asexual reproduction), but it can also mate with another cell and form a zygospore that has desiccation tolerance (sexual reproduction: Fig. 1B). As we can easily induce sexual reproduction and obtain F_1_ populations in laboratory experiments (Kawaguchi et al. 2023), we observed population dynamics of the alga under multiple stressors: salinity and acidity (low pH). Freshwater green algae go extinct under salinity stress (Lachapelle and Bell 2012, Petkovic and Colegrave 2019, 2023), and some species of the genus *Closterium* has been reported to prefer neutral or high pH water (Gough 1977, Renstrom 1979, Greenwood and Lowe 2006). By comparing parental and hybrid F_1_ populations, we observed large variation in population growth rates of F_1_ populations under low pH. This can be seen as an example of transgressive segregation, the formation of extreme phenotypes in hybrid populations (Rieseberg et al. 1999). Interestingly, even under severe environmental conditions when parental populations had negative growth rates, some F_1_ populations showed positive growth rates. This suggests the potential importance of sexual reproduction for producing genetic variation within populations that can prevent population extinction in deteriorating environments. We further found greater enrichment of genes with copy number variations in terms related to pH stress than those to salt stress in a gene ontology (GO) enrichment analysis, and discuss the potential importance of gene copy number variation and recombination for producing genetic variation based on recently sequenced genomic information for the alga (Kawaguchi et al. 2023).

**Fig. 1.**
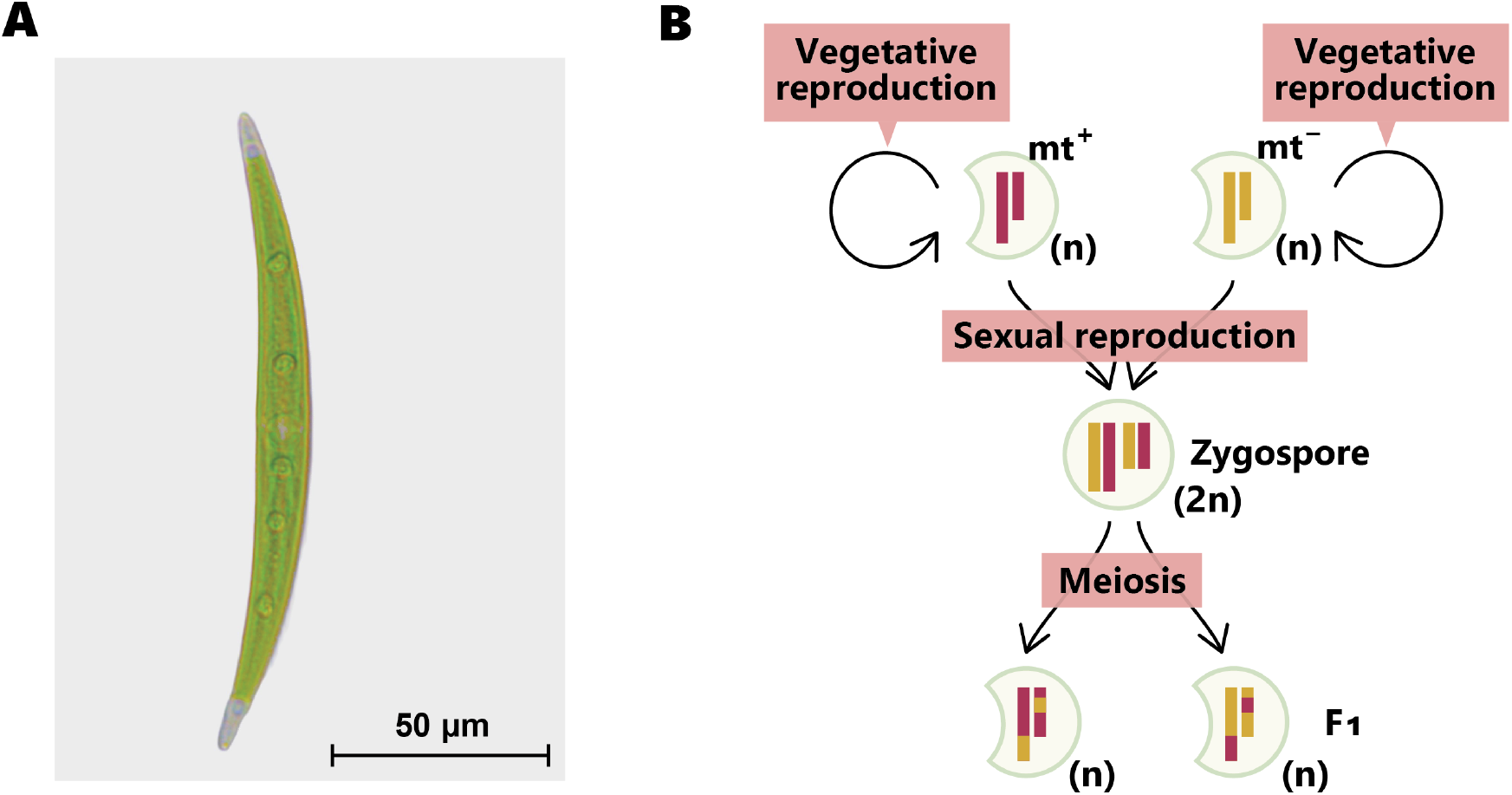
Green alga, *Closterium peracerosum–strigosum–littorale* complex, and its life cycle. (A) A cell of *C. psl*. complex (NIES-65). (B) The life cycle of *C. psl*. complex, with mating types “mt^+^” (mating type plus) and “mt^−^” (mating type minus). Chromosomes from parental strains, mt^−^ and mt^+^, are colored by orange and red, respectively. The symbols “n” and “2n” denote haploid and diploid cells, respectively. Haploid cells proliferate via vegetative reproduction, whereas sexual reproduction between different mating types produces diploid zygospores. Then, haploid cells (F_1_) were produced from the diploid zygospores through meiosis.

## Materials and Methods

### Base Population and Culture Conditions

Two strains of *C. psl*. complex, NIES-64 and NIES-65, were obtained from the National Institute for Environmental Studies (NIES), Environment Agency (Ibaraki, Japan). The strains are genetically close and sympatric (Kawaguchi et al. 2023). They are heterothallic strains and they can form zygospores between cells of two mating types: mating type plus (mt^+^: NIES-65) and mating type minus (mt^−^: NIES-64) (Tsuchikane and Sekimoto 2019, Kawaguchi et al. 2023, Ohtaka and Sekimoto 2023). They recognize each other through the sex pheromones and then form zygotes (Fig. 1B: Sekimoto et al. 1994, Nojiri et al. 1995). Through sexual reproduction, haploid parental populations produce diploid zygospores, and then haploid offspring (F_1_) populations arise via meiosis and recombination (Fig. 1B).

The base population used for the experiments was cultured in a nitrogen-supplemented medium (CA medium). The CA medium contains the following reagents in 99.9 mL of distilled water: 2 mg of calcium nitrate tetrahydrate (Ca(NO_3_)_2_ · 4H_2_O), 10 mg of potassium nitrate (KNO_3_), 5 mg of ammonium nitrate (NH_4_NO_3_), 3 mg of sodium beta-glycerophosphate pentahydrate (β-Na2glycerophosphate · 5H_2_O), 2 mg of magnesium sulfate heptahydrate (MgSO_4_ · 7H_2_O), 0.01 µg of vitamin B_12_, 0.01 µg of biotin, 1 µg of thiamine HCl, 0.1 mL of PIV metal solution, 0.1 mL of Fe (as EDTA; 1:1 molar), and 40 mg of HEPES. The PIV metal solution contains the following reagents in 100 mL of distilled water: 100 mg Na_2_EDTA · 2H_2_O, 19.6 mg FeCl_3_·6H_2_O, 3.6 mg MnCl_2_·4H_2_O, 1.04 mg ZnCl_2_, 0.4 mg CoCl_2_·6H_2_O and 0.25 mg Na_2_MoO_4_·2H_2_O (pH 7.2, 0% NaCl; Kasai et al. 2004). The culture conditions were maintained at 22 °C under a 14 h light/10 h dark cycle. The light intensity was about 40 µmol photons m^-2^ s^-1^.

### Establishment of F_1_ Populations

We grew cells from the parental strains, NIES-64 and NIES-65, vegetably for 10 days and then they were washed two times with a nitrogen-depleted mating-inducing medium (MI medium; Ichimura 1971). The MI medium contains the following reagents in 99.7 mL of distilled water: 4 mg of MgSO_4_ · 7H_2_O, 5 mg of β-Na_2_glycerophosphate · 5H_2_O, 10 mg of CaCl_2_ · 2H_2_O, 0.01 µg of vitamin B_12_, 0.01 µg of Biotin, 1 µg of thiamine HCl, 0.3 mL of PIV metals, and 50 mg of Tris (hydroxymethyl) aminomethane. Subsequently, the two strains were mixed and incubated in the MI medium at a concentration of 10,000 cells/ml for each strain. This mixture was exposed to 24 h continuous light for 10 days. The zygotes were then kept in the dark conditions for a week until they were completely dried out (note that they have desiccation tolerance). After a week, we added the CA medium to induce germination of the zygospores. Through these processes, we established eight F_1_ populations in total. Populations F_1_-A through F_1_-E germinated from 1,000 zygospores each, whereas F_1_-F germinated from 100 zygospores and F_1_-G and F_1_-H germinated from 10,000 zygospores.

### Culturing of Populations

The populations of NIES-64, NIES-65, and eight F_1_ populations were cultured in six-well plates at a density of 2,500 cells per 5 mL CA medium (Fig. S1). The culture conditions varied across combinations of NaCl concentrations (0, 0. 05, 0.1, and 0.5%) and pH levels (6.2, 7.2, and 8.2), resulting in a total of 12 different conditions. For reference, 0.5% NaCl corresponds to 5g/L NaCl and ca. 0.0856 M whereas seawater is about 3% NaCl (ca. 0.5 M). Each condition was replicated five times.

### Measurement of Growth Rate

To calculate the population growth rate, algal biomass was measured on the first day (day 0; the day the cells were introduced to each medium), as well as on days 1, 2, and 6, using a Multiskan SkyHigh TCD (Thermo) at an absorbance of 684 nm as like previous experimental studies on *Chlamydomonas* (Lachapelle and Bell 2012, Petkovic and Colegrave 2019, 2023). As the pigment content variation of a cell may bias the estimation of algal biomass, we used a wavelength where absorbance by the pigment is small (Griffiths et al. 2011). We also checked the correlation between the absorbance and the number of cells (Fig. S2). Before every measurement, the cells were agitated by pipetting. To eliminate the effects of the background, the absorbance values of media without cells were subtracted from all measurements.

The biomass dynamics was modeled using nonlinear least-squares fitting with the minpack.lm package (Elzhov et al. 2016) in R (version 4.3.2) (R Core Team 2023) as follows:

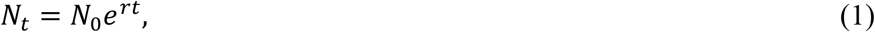

where *N*_*t*_ represents the algal biomass (absorbance values) on day *t, r* is the biomass growth rate, *t* denotes the number of days, and *N*_0_ is the algal biomass on day 0. Hereafter we use “population growth” to represent biomass growth dynamics (Fig. S3).

To investigate whether the variation in growth rates in the F_1_ populations is significantly larger than those expected from the parental (NIES-64 and NIES-65) populations, we calculated a mean and a standard deviation (SD) in population growth rates of the two parental populations. Then we sampled F_1_ individuals from a normal distribution with the calculated mean and SD. This is an expected distribution of F_1_ growth rates, and we calculated a SD of the distribution. The simulation was repeated 10,000 times to construct a null distribution of the SDs. We then examined whether the observed SDs of the F_1_ population significantly deviated from these simulated distributions (whether *p* < 0.01 or not).

## Results

We first analyzed how salinity and low pH affect population growth rates, *r*. To simplify our analyses, we focused on conditions with 0.05%, 0.1%, and 0.5% NaCl and pH 6.2 and 7.2 in the main text (Fig. 2). See Fig. S3 and S4 for the entire dataset including conditions with 0% NaCl and/or pH 8.2. We found that both stresses decreased population growth rates significantly (two-way ANOVA: salinity; *F*_2,291_ = 101, *p* < 0.001, pH; *F*_1,291_ = 46.5, *p* < 0.001, Fig. 2, Table S1). In addition, the effects of stressors were not additive (salinity × pH interaction; *F*_2,291_ = 7.24, *p* < 0.001, Fig. 2, Table S1), indicating the importance of considering two stressors simultaneously. The results from Fig. 2 were consistent with those based on the entire dataset (Table S2). As expected, we found that combining two types of stressors can further decrease population growth rates: for example, most populations showed positive growth rates in the condition with 0.05% NaCl and pH 6.2 (Fig. 2A, S4B) as well as that with 0.1% NaCl and pH 7.2 (Fig. 2E, S4G), but many of them showed negative growth rates in the condition with 0.1% NaCl and pH 6.2 (Fig. 2B, S4C). Growth responses of the two parental populations to stressors can be sometimes similar (Fig. 2B-C), but sometimes different (Fig. 2E). Under the 0.1% NaCl and pH 7.2 condition (Fig. 2E, S4G), the NIES-64 strain showed a significantly higher growth rate than the NIES-65 strain (Student’s *t*-test: *t*-value = 3.74, d.f. = 7, *p* = 0.00726).

**Fig. 2.**
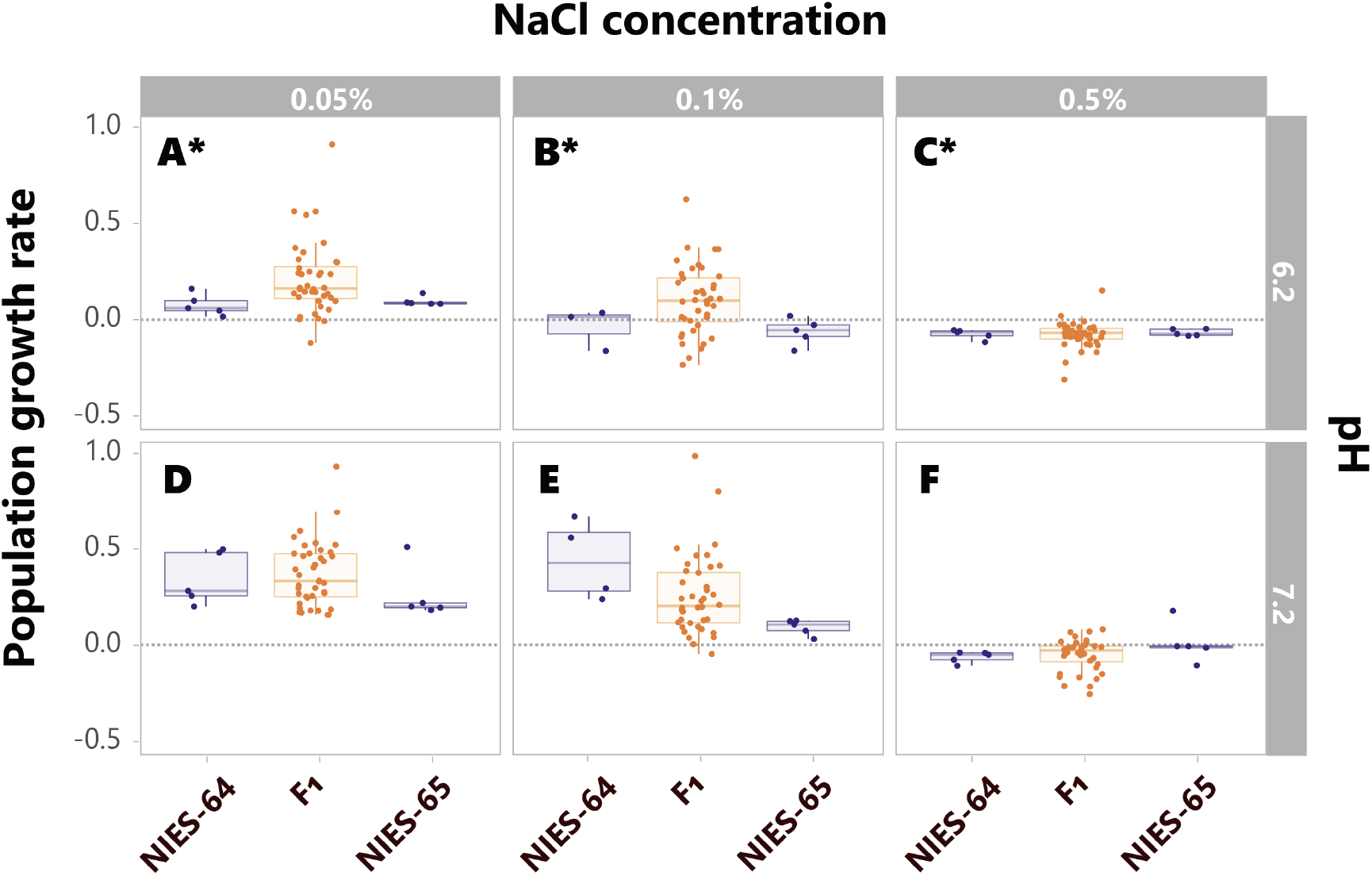
Population growth rates under each stress condition. Blue and orange points (and boxplots) denote parental (NIES-64 and NIES-65) and F_1_ populations, respectively. Bars and boxes represent the median and interquartile range, respectively. Whiskers extend to 1.5 times the interquartile range. A, D: 0.05% NaCl. B, E: 0.1% NaCl. C, F: 0.5% NaCl. A-C: pH 6.2. D-F: pH 7.2. Under condition marked with an asterisk (A, B, C), the F_1_ populations showed significantly larger variation in growth rates than the parental ones (*p* < 0.01). See Fig. S3 and S4 for more details and Fig. S5 for the effects of the founder population size.

Under the stressful conditions, some F_1_ populations exhibited larger variation in growth rates compared with the parental ones (Fig. 2). Specifically, under conditions of pH 6.2 with 0%, 0.05%, 0.1%, and 0.5% NaCl; pH 7.2 with 0%; and pH 8.2 with 0.5% NaCl, the F_1_ populations showed significantly greater standard deviations in growth rates than those expected from parental populations (*p* < 0.01, Fig. 2A-C and S4A-E and L, see Materials and Methods section for details). For example, in the condition with 0.05% NaCl and pH 6.2, the two parental populations showed very similar positive growth rates whereas the F_1_ populations exhibited large variation of growth rates: the variation is so large that some of them are even negative (Fig. 2A, S4B). Furthermore, some F_1_ populations demonstrated positive growth rates even under severe stress conditions where the parental populations showed negative growth rates (0.1% NaCl and pH 6.2, Fig. 2B, S4C). Even in the severest condition (0.5% NaCl and pH 6.2), some F_1_ populations showed positive growth rates (Fig. 2C, S4D). We also found that F_1_ populations germinated from larger number of zygospores exhibited higher growth rates in two conditions (Fig. S5D, S5F), but it is not always the case (see Fig. S5 for details).

## Discussion

Sexual reproduction has been a central enigma in evolutionary biology because of the many costs of sex (Williams 1975, Maynard Smith 1978, Bell 1982, Hartfield and Keightley 2012, Lehtonen et al. 2012). Recent studies have shown that sexual reproduction is important for organisms to adapt to rapidly changing environments (Colegrave 2002, Kaltz and Bell 2002, Goddard et al. 2005, Morran et al. 2009, Becks and Agrawal 2010, Gray and Goddard 2012) and for preventing population extinction (Lachapelle and Bell 2012, Petkovic and Colegrave 2019, 2023), but it was not clear whether a single episode of sexual reproduction can produce large amount of genetic variation. By using the unicellular green alga, *Closterium peracerosum–strigosum– littorale* complex, we have shown large variation in population growth rates of F_1_ populations (Fig. 2, S4). Even when parental populations show negative growth rates, some F_1_ populations show positive growth rates in severe environmental conditions (e.g., the condition with 0.1% NaCl and pH 6.2, Fig. 2B, S4C). Our results indicate that, although asexual reproduction is more efficient in population growth, sexual reproduction is important for growth in stressful environments and for evolutionary rescue.

Theory suggests that recombination due to sexual reproduction has two antagonistic effects on evolutionary rescue: it generates and breaks up favorable gene combinations (Uecker and Hermisson 2016, Orive et al. 2017). The former is related to the Fisher-Muller hypothesis where sexual recombination can combine beneficial alleles into the same genome (Fisher 1930, Muller 1932). Our results are consistent with the two classical ideas: sexual reproduction tended to increase the variance in population growth rates, and it resulted in positive (or negative) growth of some F_1_ populations even when their parental populations showed negative (or positive) growth (Fig. 2, S4). Genome information recently obtained for this algal species (*C. psl*. complex) has revealed that approximately 30% of the genes in the parental strains showed copy number differentiation (Kawaguchi et al. 2023). Variation in the number of gene copies may produce large variation in the F_1_ generation through recombination (Hastings et al. 2009). Thus, it will be interesting to carefully consider how genetic architecture, including structural variation, affects evolutionary rescue in future studies (Gomulkiewicz et al. 2010, Yamamichi 2022).

When we regard population growth rate as a quantitative trait, it is possible to consider our results as an example of transgressive segregation (Rieseberg et al. 1999). Previous studies tended to focus on its effects on macroevolution such as speciation and adaptive radiation (e.g., Kagawa and Takimoto 2018), but our results suggest that it may affect rapid contemporary evolution as well as evolutionary rescue. Kawaguchi et al. (2023) estimated the genome sizes of the F_1_ populations between NIES-64 and NIES-65 and found that they were intermediate values between those of their parental populations (Fig. 3a of Kawaguchi et al. 2023). This is consistent with the results on population growth rates in, for example, Fig. 2E, but not consistent with the results under low pH (Fig. 2A-C). Thus, it will be important to explore causes of the difference between the estimated genome size and population growth rates in future studies. In this context, it will be helpful to conduct linkage mapping to understand which genes are responsible for population growth rates. Green algae are suitable as they grow as haploids, and thus it should be easier to detect genes responsible for a quantitative trait without considering allelic dominance.

We observed that the green alga grew very well in high pH medium (when pH was 8.2, Fig. S3, S4), which is consistent with previous studies on some species of the genus *Closterium* (Gough 1977, Renstrom 1979, Greenwood and Lowe 2006). On the other hand, low pH reduced growth rates significantly as expected (when pH was 6.2, Fig. 2, S3, S4). Interestingly, the F_1_ populations tended to have larger variation in growth rates in low pH environments (Fig. 2A-C). This may be seen as an example of cryptic genetic variation, which has little or no effect on phenotype normally but generates phenotypic variation under novel conditions (Paaby and Rockman 2014).

Previous studies on *Chlamydomonas* species have suggested that genes encoding heat-shock proteins and plasma membrane H^+^-ATPase were attributable to adaptation to low pH environments (Gerloff-Elias et al. 2006, Hirooka et al. 2017). In the genus *Klebsormidium*, belonging to streptophyte algae (as like *Closterium*), it has been demonstrated that the ability to adapt to low pH conditions has evolved multiple times independently (Škaloud et al. 2014). These previous studies, combined with our findings, imply that adaptation to low pH environments may be related to cryptic genetic variation that can promote rapid adaptation.

Under high salt stress, we also observed a notable reduction in population growth rate (Fig. 2, S3, S4). Salt is known to impose osmotic and oxidative stresses by disrupting the homeostasis of ions and inhibits growth by decreasing the rates of photosynthesis (Husic and Tolbert 1986, Neelam and Subramanyam 2013) and promote population extinction in *Chlamydomonas* (Lachapelle and Bell 2012, Petkovic and Colegrave 2019, 2023). However, unlike low pH, salt stress alone did not result in increased variation in growth rates of F_1_ populations (Fig. 2D-F). Our additional analyses suggest that the large variation observed in the F_1_ populations under low pH may be attributed to copy number variation between parental strains. Previous studies have reported that there was a difference in chromosome numbers between the parental strains NIES-64 and NIES-65, and approximately 30% of genes exhibited variations in copy number (Kawaguchi et al. 2023, Tsuchikane et al. in press). Following the methodology of Kawaguchi et al. (2023), our Gene Ontology (GO) enrichment analysis revealed greater enrichment of genes with copy number variations in terms related to pH stress than those to salt stress (see Fig. S6 for more details). Notably, the term “regulation of pH” (GO:0006885) showed significant enrichment (*p* = 0.0339). It suggests that the copy number variation contributes to the greater variance observed in F_1_ populations under low pH conditions. Further analysis comparing genes involving in pH and salt stress responses would be valuable, providing insights into how genetic structures, environmental stressors, and recombination contribute to transgressive segregation and promote evolutionary rescue.

While previous studies tended to focus on a single environmental factor to understand its deteriorating effect on population growth and evolutionary rescue (Kinnison and Hairston 2007, Alexander et al. 2014, Carlson et al. 2014, Bell 2017), recent studies proposed that multiple stressors are simultaneously affecting ecosystem dynamics in the Anthropocene (Orr et al. 2020). Therefore, it will be important to consider interacting multiple stressors in future studies on evolutionary rescue to deepen our understanding on conservation of endangered species as well as management of target species for hunting and fisheries. For archiving this goal, it will be essential to utilize genome information to reveal the genetic architecture underlying adaptation. If there are two or more stresses, more loci will affect phenotypic adaptation to the degraded environments, and it will further make sexual recombination important for producing combinations better adapted to the environmental changes.

## Author contributions

Conceptualization: MY and YWK; Experiments and data analyses: YWK; Funding acquisition: MY; Writing-original draft: MY; Writing-reviewing and editing: MY and YWK.

## Acknowledgments

*C. psl*. complex (NIES-64 and NIES-65) were provided by NIES through the National BioResource Project (NBRP) of the Ministry of Education, Culture, Sports, Science and Technology (MEXT), Japan. We thank T. Tsuchimatsu, M. Aono, S. Shibasaki, and N. G. Hairston Jr. for their helpful comments and laboratory works.

