## Supplementary materials for "A single episode of sexual reproduction can prevent population extinction under multiple stressors"

**Supplementary Table S1. The effects of salinity and pH conditions on the population growth rates.**

The output of analysis of variance (two-way ANOVA) for population dynamics under two stressors (0.05%, 0.1%, and 0.5% NaCl and pH 6.2 and 7.2). The reference categories for NaCl and pH were set to 0.05% NaCl and pH 6.2, respectively.

| | **Df** | **Sum Sq** | **Mean Sq** | ***F* value** | **Pr(>*F*)** |
|---|---|---|---|---|---|
| NaCl | 2 | 5.64 | 2.82 | 101 | 3.97e-34 |
| pH | 1 | 1.29 | 1.30 | 46.5 | 5.39e-11 |
| NaCl × pH | 2 | 0.402 | 0.201 | 7.24 | 8.58e-04 |
| Residuals | 291 | 8.10 | 0.0278 | | |

**Supplementary Table S2. The effects of salinity and pH conditions on the population growth rates including all data.**

The output of analysis of variance (two-way ANOVA) for population dynamics under two stressors including all data (0%, 0.05%, 0.1%, and 0.5% NaCl and pH 6.2, 7.2, and 8.2). The reference categories for NaCl and pH were set to 0% NaCl and pH 6.2, respectively.

| | **Df** | **Sum Sq** | **Mean Sq** | ***F* value** | **Pr(>*F*)** |
|---|---|---|---|---|---|
| NaCl | 3 | 17.0 | 5.68 | 208 | 5.69e-92 |
| pH | 2 | 5.74 | 2.87 | 105 | 9.03e-40 |
| NaCl × pH | 6 | 2.53 | 0.421 | 15.4 | 1.76e-16 |
| Residuals | 585 | 15.9 | 0.0273 | | |

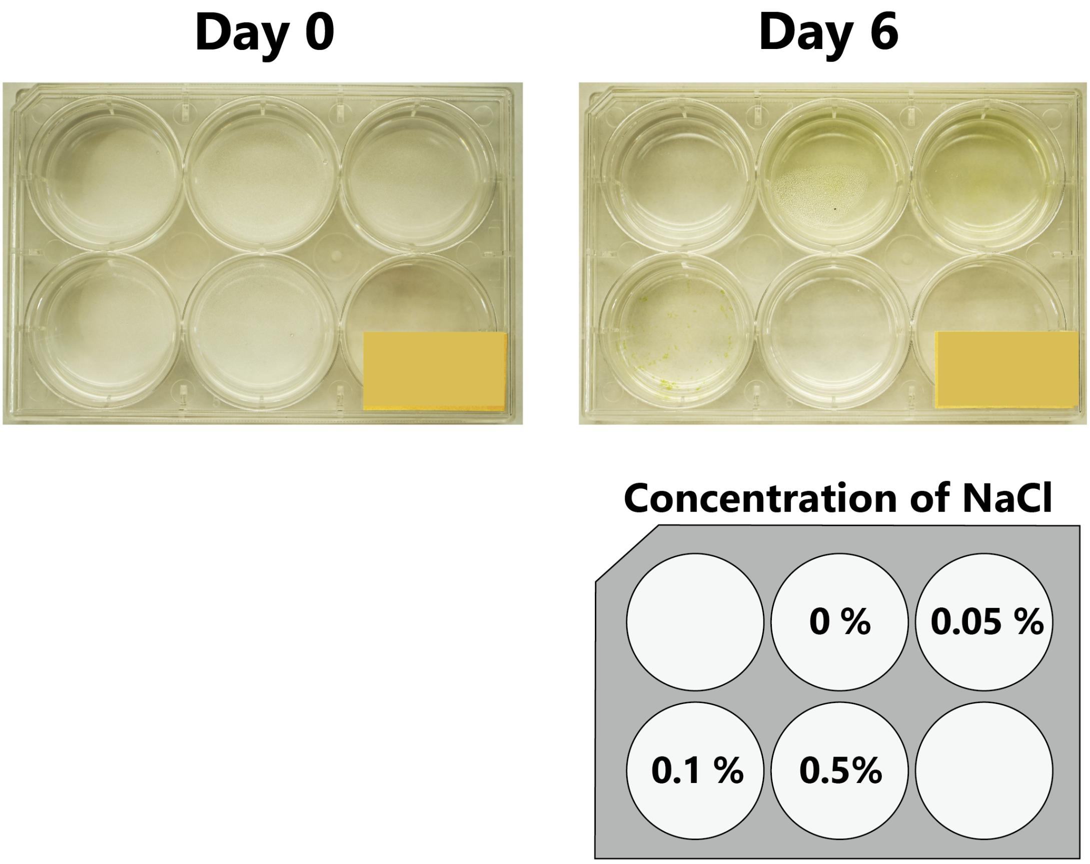


**Supplementary Figure S1. An example of population dynamics.**

Photographs of the $F_1$-C population at pH 7.2 on Day 0 and Day 6. A bottom-right diagram indicates the salinity concentration in each well. The four cells were almost transparent on Day 0 and some of them had green cells (e.g., 0%, 0.05%, and 0.1% NaCl) on Day 6. As high salinity reduced the population growth, the cell of 0.5% NaCl was almost transparent.

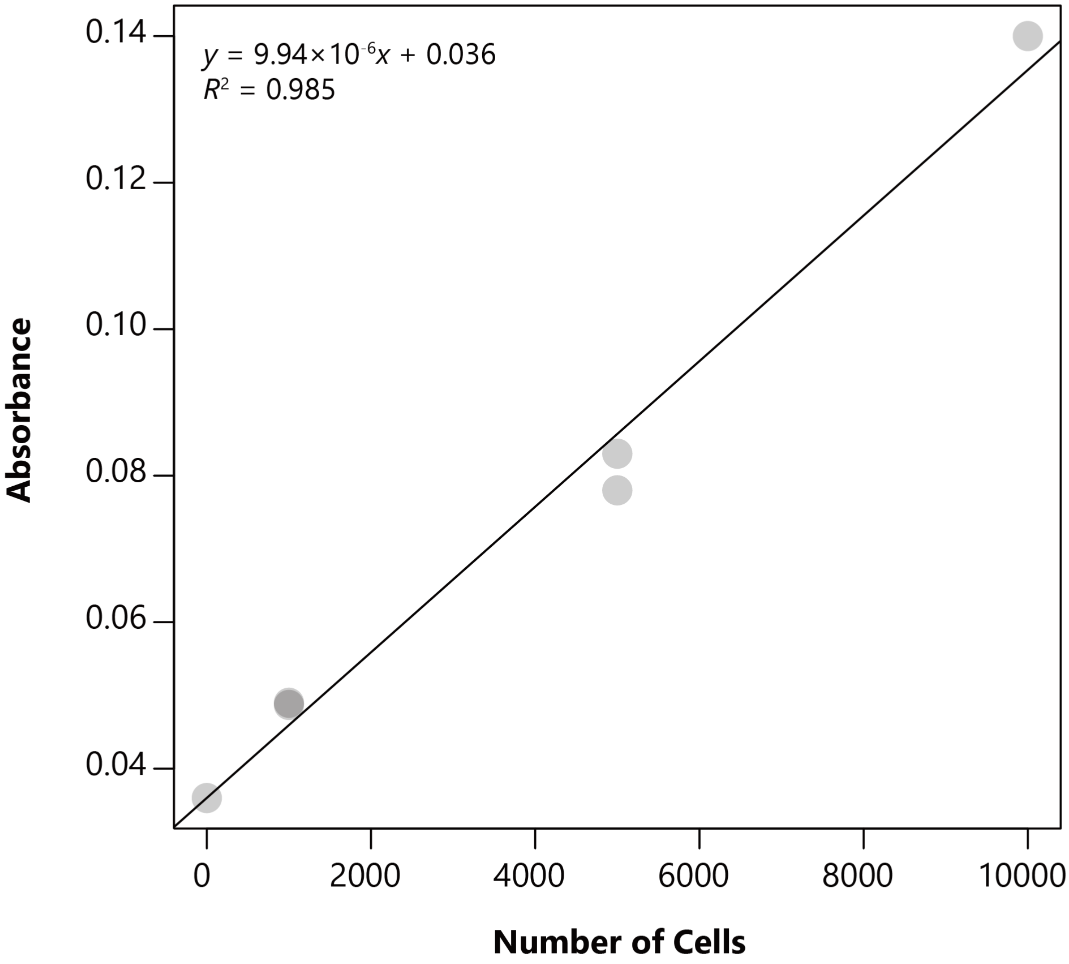


**Supplementary Figure S2. Calibration curve between the number of cells and the absorbance.**

Relationship between the number of cells and the absorbance at 684 nm when pH was 7.2 and NaCl was 0%. A solid line represents a result of linear regression ($p = 8.09 \times 10^{-5}$).

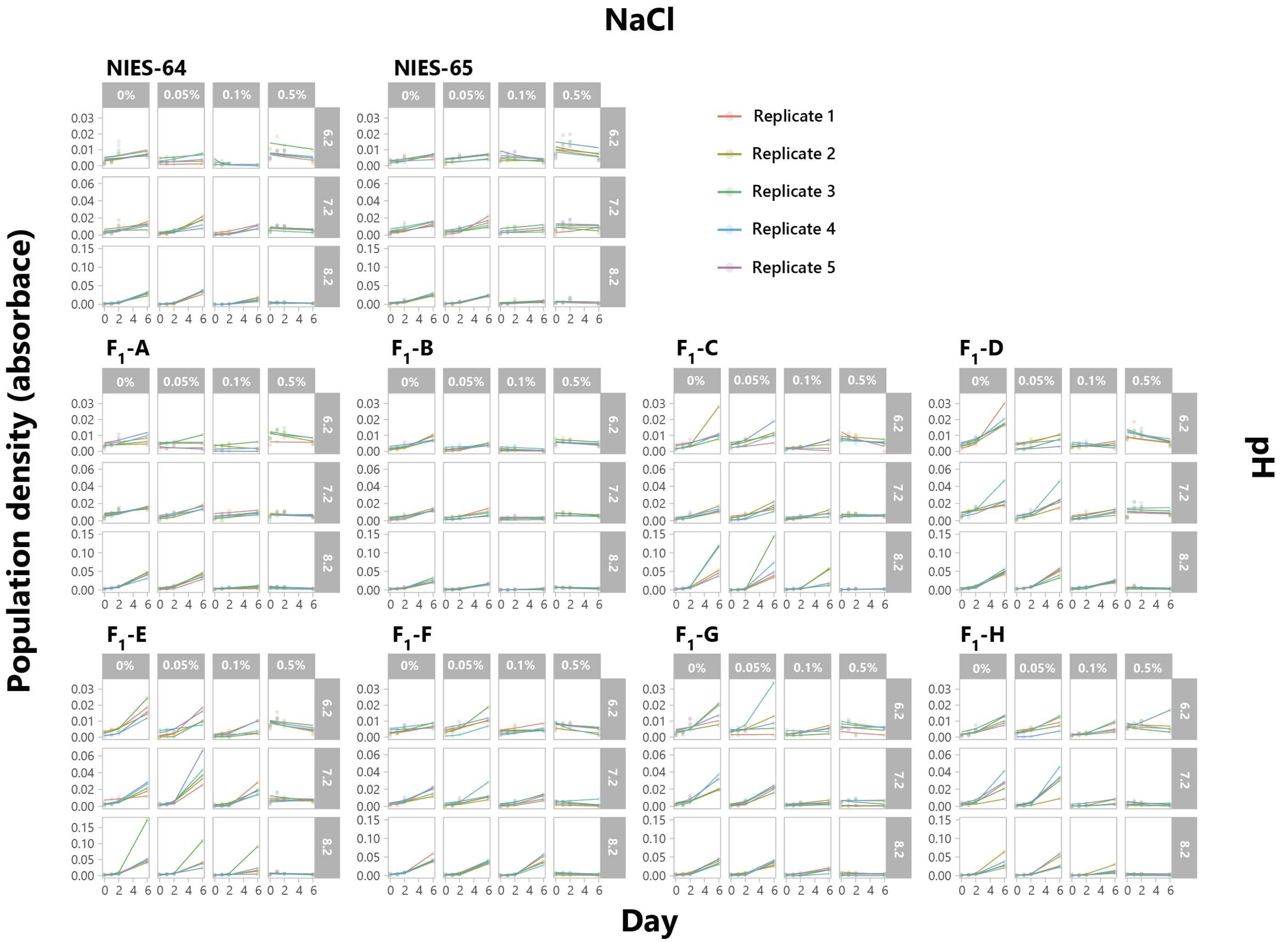


**Supplementary Figure S3. Population dynamics and fitted curves under various conditions.**

Data points were obtained at Day 0, 1, 2, and 6. Solid lines represent the fitted curves obtained using a nonlinear least-squares fitting model (equation 1 in the main text). Each color represents five replicates. NIES-64 and NIES-65 are parental populations and $F_1$-A to

$F_1$-H populations are eight hybrid $F_1$ populations. Note that the maximum values of Y-axis are different depending on pH.

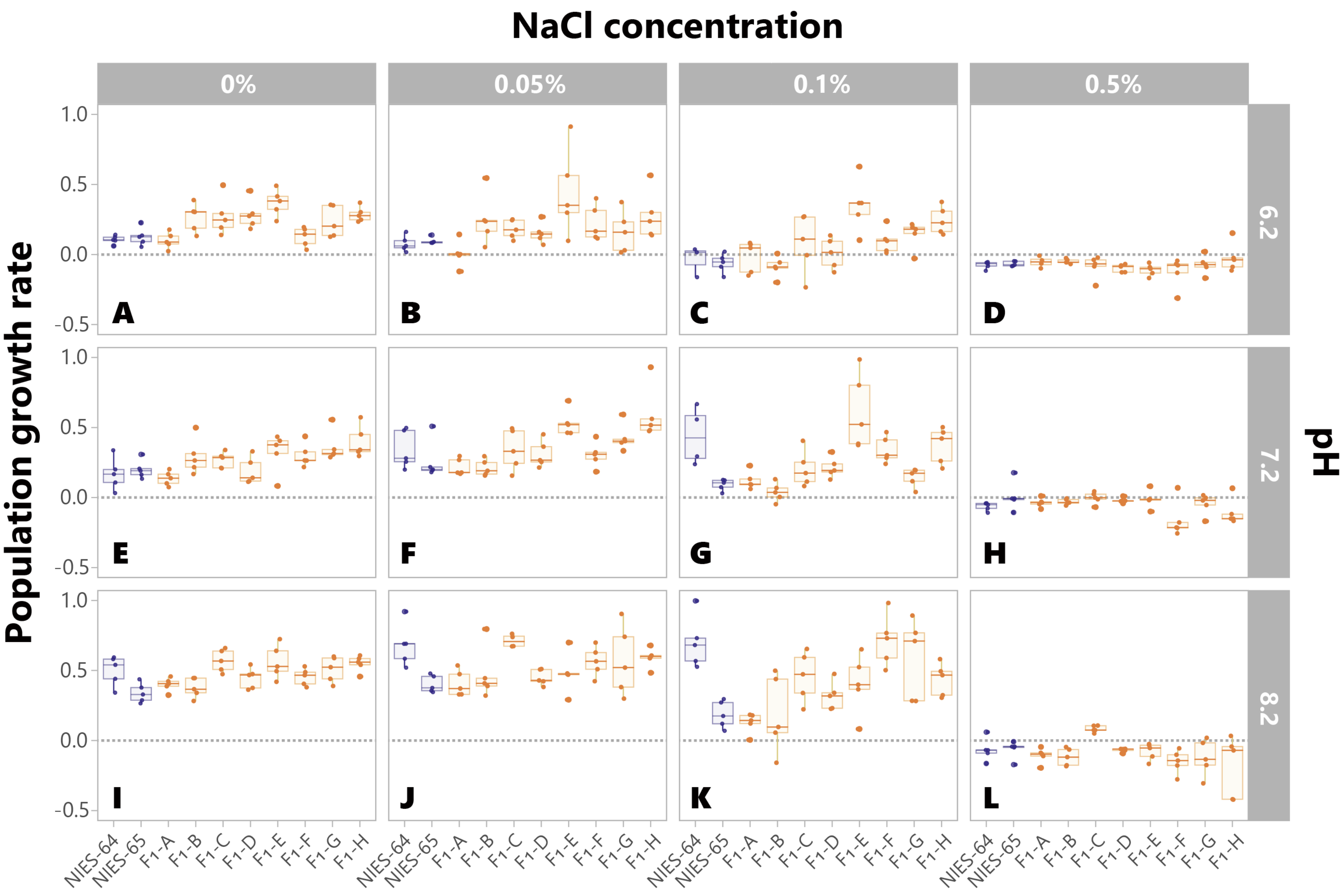


**Supplementary Figure S4. Population growth rates under various conditions.**

Blue and orange denote the parental (NIES-64 and NIES-65) and $F_1$ populations ($F_1$-A to $F_1$-H), respectively. Bars and boxes represent the medians and the interquartile ranges, respectively. Whiskers extend to 1.5 times the interquartile ranges. Gray dashed lines indicate growth rate 0. In panels A-E and L, the $F_1$ populations exhibited significantly greater standard deviations in growth rates than the parental populations ($p < 0.01$).

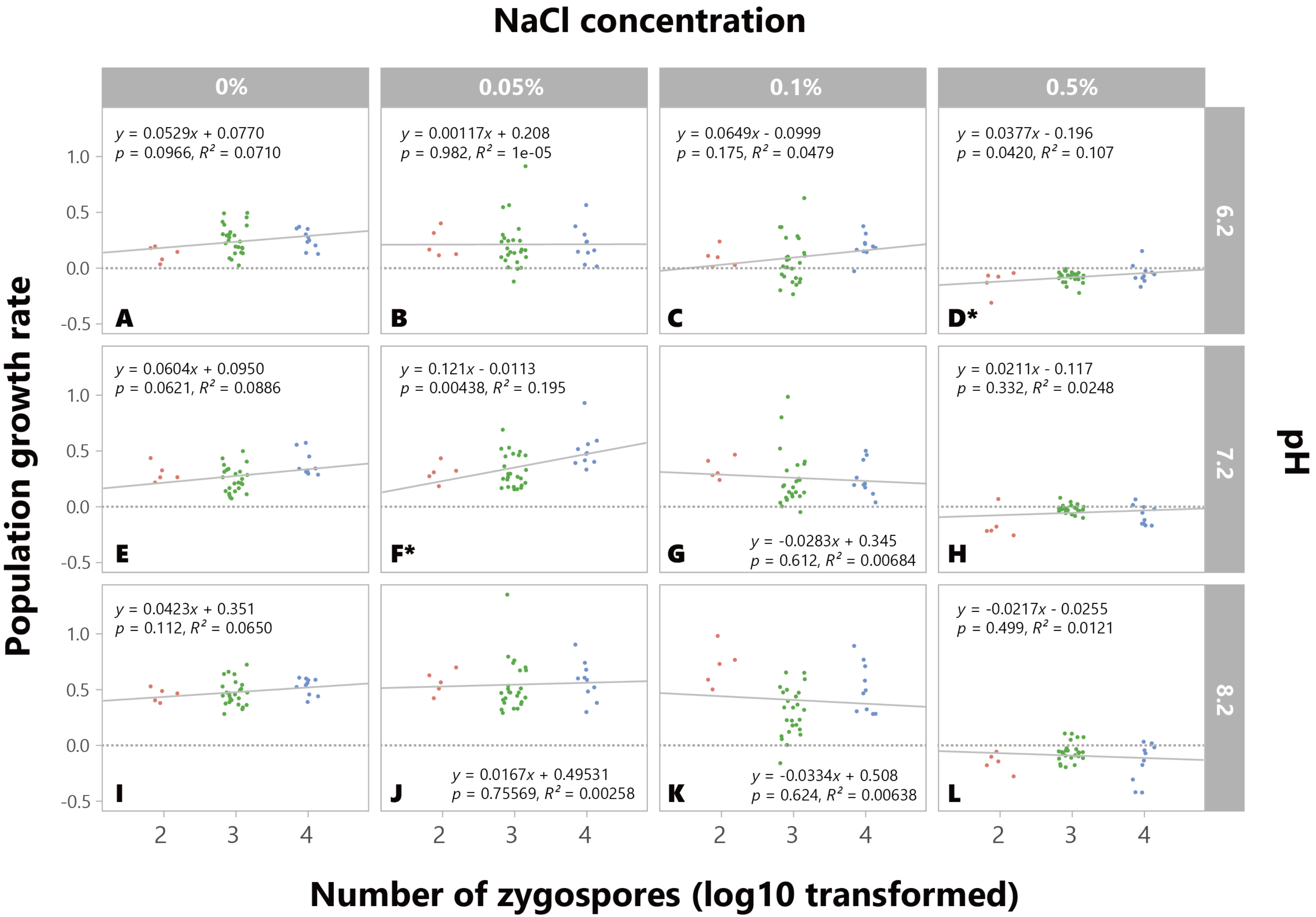
NaCl concentration
0%
0.05%
0.1%
0.5%
6.2
7.2
8.2
pH
Population growth rate
Number of zygospores (log10 transformed)
y = 0.0529x + 0.0770
p = 0.0966, R² = 0.0710
A
y = 0.00117x + 0.208
p = 0.982, R² = 1e-05
B
y = 0.0649x - 0.0999
p = 0.175, R² = 0.0479
C
y = 0.0377x - 0.196
p = 0.0420, R² = 0.107
D*
y = 0.0604x + 0.0950
p = 0.0621, R² = 0.0886
E
y = 0.121x - 0.0113
p = 0.00438, R² = 0.195
F*
y = -0.0283x + 0.345
p = 0.612, R² = 0.00684
G
y = 0.0211x - 0.117
p = 0.332, R² = 0.0248
H
y = 0.0423x + 0.351
p = 0.112, R² = 0.0650
I
y = 0.0167x + 0.49531
p = 0.75569, R² = 0.00258
J
y = -0.0334x + 0.508
p = 0.624, R² = 0.00638
K
y = -0.0217x - 0.0255
p = 0.499, R² = 0.0121
L

55

**Supplementary Figure S5. The effects of the number of zygospores on the population growth rate.**

Among the established eight $F_1$ populations, one population germinated from 100 zygospores (orange), five populations germinated from 1,000 zygospores (green), and one population germinated from 10,000 zygospores (blue). Gray dashed lines indicate growth rate 0. We performed linear regressions to analyze the relationship between the number of zygospores and their growth rates. For these analyses, we applied a log10 transformation to the independent variable, which is the number of zygospores. Their results are shown as gray solid lines and as formulas on each panel. Under condition marked with an asterisk (D, F), their growth rates were significantly correlated with their number of zygospores ($p < 0.05$, D; $t$ =2.11, 95% confidence interval of slope = 0.00145 to 0.0740, F; $t$ = 3.03, 95% confidence interval of slope = 0.0402 to 0.202). However, when considering multiple testing, we cannot say that there was a clear correlation between the growth rate and number of zygospores.

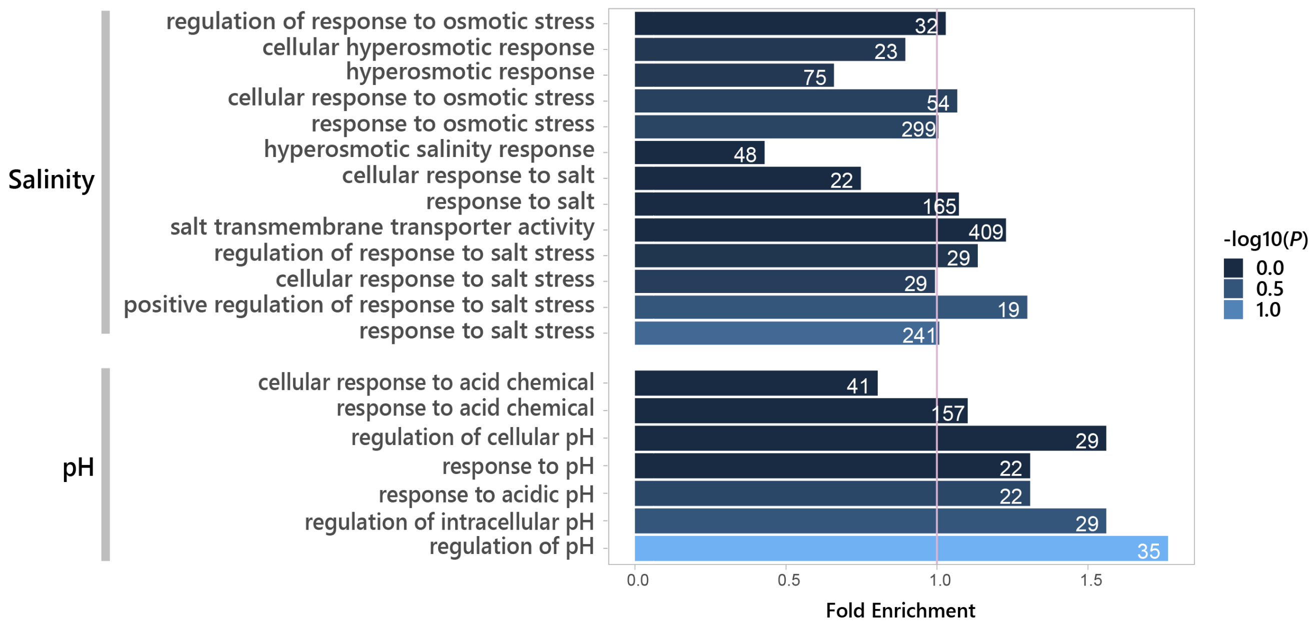


**Supplementary Figure S6. GO analysis of functions with response to salinity and pH stress.** We conducted a Gene Ontology (GO) enrichment analysis of genes with copy number variation between parental strains (NIES-64 and NIES-65). Following the methodology of Kawaguchi et al. (2023), we counted the copy number for each gene from NIES-64 and NIES-65 (accession number: GCA_948144035.1 and GCA_948144175.1) and assigned their respective GO terms. The enrichment analysis was performed using topGO with the “weight01” algorithm (Alexa et al., 2006) and GO.db v.3.12.1 (Carlson et al., 2019) in R (version 4.3.2) (R Core Team 2023). This figure shows the result for terms related to salinity and pH stress responses, with the number of genes annotated for each GO term shown on the bars. Notably, the term “regulation of pH” (GO:0006885) was significantly enriched in genes with copy number variation ($p = 0.0339$), The terms related to pH stress responses exhibiting higher enrichment than estimated from the overall gene copy number pattern (fold enrichment > 1), indicating that genes associated with pH stress response tend to exhibit copy number variation. It suggests that the copy number variation might contribute to the greater variance observed in $F_1$ populations under low pH conditions.

R Core Team. 2023. R: A language and environment for statistical computing. R Foundation for Statistical Computing, Vienna, Austria.
